# CHAPERON*g*: A tool for automated GROMACS-based molecular dynamics simulations and trajectory analyses

**DOI:** 10.1101/2023.07.01.546945

**Authors:** Abeeb Abiodun Yekeen, Olanrewaju Ayodeji Durojaye, Mukhtar Oluwaseun Idris, Hamdalat Folake Muritala, Rotimi Olusanya Arise

## Abstract

Molecular dynamics (MD) simulation is a powerful computational tool used in biomolecular studies to investigate the dynamics, energetics, and interactions of a wide range of biological systems at the atomic level. GROMACS is a widely used free and open-source biomolecular MD simulation software recognized for its efficiency, accuracy, and extensive range of simulation options. However, the complexity of setting up, running, and analyzing MD simulations for diverse systems often poses a significant challenge, requiring considerable time, effort, and expertise. Here, we introduce CHAPERON*g*, a tool that automates the GROMACS MD simulation pipelines for protein and protein-ligand systems. CHAPERON*g* also integrates seamlessly with GROMACS modules and third-party tools to provide comprehensive analyses of MD simulation trajectories, offering up to 20 post-simulation processing and trajectory analyses. It also streamlines and automates established pipelines for conducting and analyzing biased MD simulations via the steered MD-umbrella sampling workflow. Thus, CHAPERON*g* makes MD simulations more accessible to beginner GROMACS users whilst empowering experts to focus on data interpretation and other less programmable aspects of MD simulation workflows.

## 1 Introduction

Molecular dynamics (MD) simulation is a robust and valuable tool for studying the dynamic behavior, energetics, and interactions of diverse biological systems, including proteins, protein-ligand complexes, nucleic acids, and membrane lipids [1]. These simulations provide insights, in full atomic details and at precise temporal resolutions, into the dynamics, stability, and functional properties of biomolecules, complementing experimental observations and providing guidance for further investigations [2, 3]. GRO-MACS [4] is a widely used MD simulation software. It is one of the gold standards for biomolecular simulation not only because of its efficiency, accuracy, and extensive range of simulation options but also because it’s a free and open-source software with a huge community of users and developers [5, 6]. MD simulation protocols typically consist of three main stages: system setup, MD production or simulation run, and trajectory analysis [7]. While advances in computational power and resources have improved the capabilities of MD simulation tools, setting up and running MD simulations with GROMACS (and other MD simulation codes) still present several challenges [8].

GROMACS is a command-line program and, as with most other popular MD simulation engines, is characterized by limited accessibility to many researchers who, despite possessing the necessary domain knowledge to interpret the relevant computational results, may not be familiar with working on the command line [5, 6]. In addition, setting up and running MD simulations requires a lot of manual tasks, making the process time-consuming, labor-intensive, and error-prone. Since many of these steps are repetitive and programmable, automation tools would greatly minimize manual user interventions, thereby improving efficiency and empowering users (beginners and experts) to focus on other aspects like parameter optimization, data analysis, and result interpretation [9].

Furthermore, MD simulations typically generate huge amounts of trajectory data, but processing the data into particularly meaningful, relevant, and informative forms often requires programming and data analysis skills [10, 11]. Thus, beginner and intermediate users with limited or no such skills are only able to gain superficial insights from MD output data in spite of MD simulation being a computation-intensive process. The ability to transform simulation data into more interpretable forms and consequently obtain optimally useful information would facilitate the gaining of relevant biological (structural and functional) insights from MD simulations [11].

Several tools integrated into standalone or web-based graphical user interfaces (GUIs) have been developed to assuage the aforementioned challenges. The earliest GUI-based programs including GUIMACS [12], jSimMacs [13], and GROMITA [14] that offered some capability to carry out GROMACS MD simulation of protein (only) systems have not been updated for a long time, making them incompatible with recent GROMACS versions [6]. Other GUI-integrated plugins like Dynamics PyMOL plugin [5, 15], Enlighten2 (a PyMOL plugin and Python package) [16], and YAMACS (a YASARA plugin) [6] have such limitations as restrictions to the simulation of specific systems (protein only or protein complexes), support for only select force field(s), lack of trajectory analysis functions, non-trivial installation of dependencies, or the need to learn other software interfaces upon which they depend [16]. MDWeb [17] and WebGro [18] are two web-based GUIs for MD simulation over limited durations. MDWeb does not support the simulation of protein-ligand complexes, while WebGro usage is limited to the GROMOS force field.

In this work, we present CHAPERON*g*, a <u>c</u>ompre<u>h</u>ensive <u>a</u>utomated <u>p</u>ip<u>e</u>line for <u>G</u>ROMACS MD simulations and t<u>r</u>aject<u>o</u>ry a<u>n</u>alyses. CHAPERON*g* is a command-line interface to GROMACS that automates and streamlines the entire MD simulation protocols for protein, protein-ligand, and protein-DNA systems. It supports ligand topology parameters obtained from popular external parametrization programs for the CHARMM, AMBER, GROMOS, and OPLS force fields. CHAPERON*g* seamlessly integrates with GROMACS modules and third-party tools to enable an extensive automated workflow of up to 20 different post-simulation trajectory and end-point analyses. In addition, it automates the steered MD and umbrella sampling simulations, a biased enhanced simulation protocol often employed to overcome sampling limitations and investigate rare events. Thus, CHAPERON*g* would not only make MD simulation more accessible to beginner GROMACS users but also expand the toolset of experts by facilitating improved efficiency and providing a platform upon which advanced and customized analyses and scripting could be built.

## 2 Methods and code implementation

CHAPERON*g* has been developed using the Bash shell scripting and the Python 3 programming language. The framework and primary modules of the CHAPERON*g* source code were written using Bash shell scripting because GROMACS is a Linux-based software. This allows a seamless GROMACS-CHAPERONg integration and ensures that the only real dependency of CHAPERON*g* is simply a functional GROMACS installation. Thus, the entire MD simulation pipelines can be automatically executed without the need for installation of additional dependencies or software save those required by GROMACS itself.

Other modules of CHAPERON*g* which provide additional and advanced functionalities are written using Python. Various Python libraries are used including Numpy [19], Pandas and Scipy [20]; for data manipulation and numerical and scientific calculations, and Matplotlib [21]; for generating graphical plots and figures. PyMOL [22], ImageMagick, or ffmpeg is used for generating simulation movies. Secondary structure elements are analyzed using DSSP [23]. Xmgrace is used for graph plotting and conversion.

The MD DaVis package [11] is used for the construction of hydrogen bond matrices and interactive three-dimensional visualizations of free energy landscapes. Installation of CHAPERON*g* is achieved by simply running the installation script provided in the package. To make all features offered by CHAPERON*g* easily accessible to users, an isolated Anaconda Python environment with all needed dependencies can be set up by running a conda setup script also provided in the package.

CHAPERON*g* offers automated GROMACS-based workflows for unbiased conventional MD simulation of protein(-only) and protein complex systems, using established and previously reported protocols [24, 25]. In addition, up to 20 automated analysis types covering system setup and simulation quality control analyses as well as post-simulation trajectory analyses are provided. A GROMACS-based workflow for the steered MD and enhanced umbrella sampling simulations for protein complexes are also automated [26, 27, 24]. Automated quality control analyses and the WHAM (weighted histogram analysis method)-based free energy calculations [28, 29] are also provided for the biased simulations.

## 3 CHAPERON*g* features and functionalities

CHAPERON*g* can be run in one of two modes of automation depending on the user’s choice; either as full-auto or semi-auto. In the full-auto mode, all simulation steps and post-simulation analyses are automatically carried out based on the simulation type and user-provided parameters. This greatly reduces repetitive and tedious manual interventions, and the user is only prompted for inputs in a few cases where automatic or pre-defined choices would not be suitable. The semi-auto mode still has most of the simulation and analyses automated, but the user is prompted more for inputs and confirmation of automatically selected choices to give more flexibility and control over the simulation parameters.

### 3.1 Automated conventional MD simulation

CHAPERON*g* offers automated MD pipelines for various systems, namely protein-only (including protein-protein complexes), protein-ligand complexes, and protein-DNA complexes. For protein-ligand systems, the pipeline recognizes small molecule ligand topologies generated via popular ligand parameterization programs and webservers, including CGenFF (for CHARMM) [30], ACPYPE (for AMBER) [31], PRODRG2 (for GROMOS) [32], and LigParGen (for OPLS) [33]. The automated protocol is organized into 12 major steps, enabling the user to start or resume from any step of the simulation process. The minimum input files required to run CHAPERON*g* are the starting structure and appropriate GROMACS .*mdp* files.

#### 3.1.1 System preparation and quality control analyses

Once launched, CHAPERON*g* automatically runs through the conversion of the input structure file to the GROMACS format, generation of protein topology (and ligand topology, if applicable), definition of the simulation box, addition of ions to the system, energy minimization and a two-stage NVT/NPT equilibration. For each of these steps, the user has full control over how the system is set up. The system and topology files are automatically updated accordingly, depending on the type of system. Following the energy minimization and equilibration steps, quality control analyses–such as the progression of the potential energy, density, temperature, pressure, and other thermodynamic parameters–are run. These enable the user to monitor the convergence of these indices and, hence, the quality of the simulation system.

#### 3.1.2 MD simulation

Following a successful setup of the system, the production MD run proceeds for the duration specified by the user in the corresponding parameter file. CHAPERON*g* also offers the option to call GROMACS to extend a previously completed run, or to resume a terminated simulation. Similar to the system preparation stage, several quality control indices including some thermodynamic parameters are analyzed and produced as Xmgrace .*xvg* files as well as publication-quality .*png* images.

### 3.2 Post-simulation processing and trajectory analyses

CHAPERON*g* provides the capability to carry out up to 20 post-simulation processing and trajectory analyses. These analyses, enabled by modules available in GROMACS, CHAPERON*g*, and other third-party tools, include root mean square deviation (RMSD), root mean square fluctuation (RMSF), radius of gyration (Rg), solvent accessible solvent area (SASA), hydrogen bond (Hbond) analysis, principal component analysis (PCA), secondary structure analysis, clustering analysis, simulation movie, two- and three-dimensional (visualizations) free energy landscapes (FELs), kernel density estimation (KDE), interactive hydrogen bond matrix, MM-PBSA (Molecular mechanics Poisson–Boltzmann surface area) free energy calculations, and multiple quality control analyses. The plots from the analyses are generated as .*xvg* and publication-quality .*png* files. These analyses provide valuable computational metrics for characterizing the stability, folding, conformational changes, interactions and dynamics of biomolecules. For example, they help in comparing different molecular dynamics simulation trajectories, analyzing the impact of mutations or ligand binding, and assessing the accuracy of the simulated models with respect to experimental data.

#### 3.2.1 RMSD, RMSF, Rg and SASA

The RMSD, RMSF, and Rg are three important structural metrics used to characterize the MD simulation of biomolecular systems [34]. The RMSD, Rg, and RMSF are computed in GROMACS by the *gmx rms, gmx gyrate*, and *gmx rmsf* modules, respectively. The RMSD measures the average distance between the atoms of a structure at an instant of the simulation against the reference starting structure. Thus, it is used to analyze the overall time-dependent structural deviation or similarity between the structures recorded in the trajectory [35, 36]. The RMSD plot of the protein (and that of the ligand in the case of a protein-ligand complex) is generated as .*xvg* files and .*png* figures.

The RMSF, like the RMSD, is a common mobility measure that quantifies the local fluctuations or flexibility within a biomolecule during simulation by measuring the average atomic or residue-level deviations [35]. The RMSF provides insights into the dynamic regions of proteins such as flexible loops, and can indicate the importance of specific residues in conformational changes or protein-ligand interactions [37]. The Rg is a commonly used measure of the compactness of protein molecules, with smaller Rg values indicating a more compact or folded structure, and larger Rg values suggesting more extended or flexible conformations during the simulation [38].

SASA is a metric that provides information about the exposed surface area of a biomolecule that is accessible to the solvent molecules. It is commonly used to investigate protein folding and stability, as well as to characterize the interaction of a protein with the surrounding solvent [39]. SASA is computed in GROMACS by the *gmx sasa* module, which employs the double cubic lattice method [40]–a variant of the “rolling ball” algorithm of Shrake and Rupley [41]. Figure 1 shows some examples of the automatically generated figures of the RMSD, RMSF, Rg, and SASA plots.

**Figure 1:**
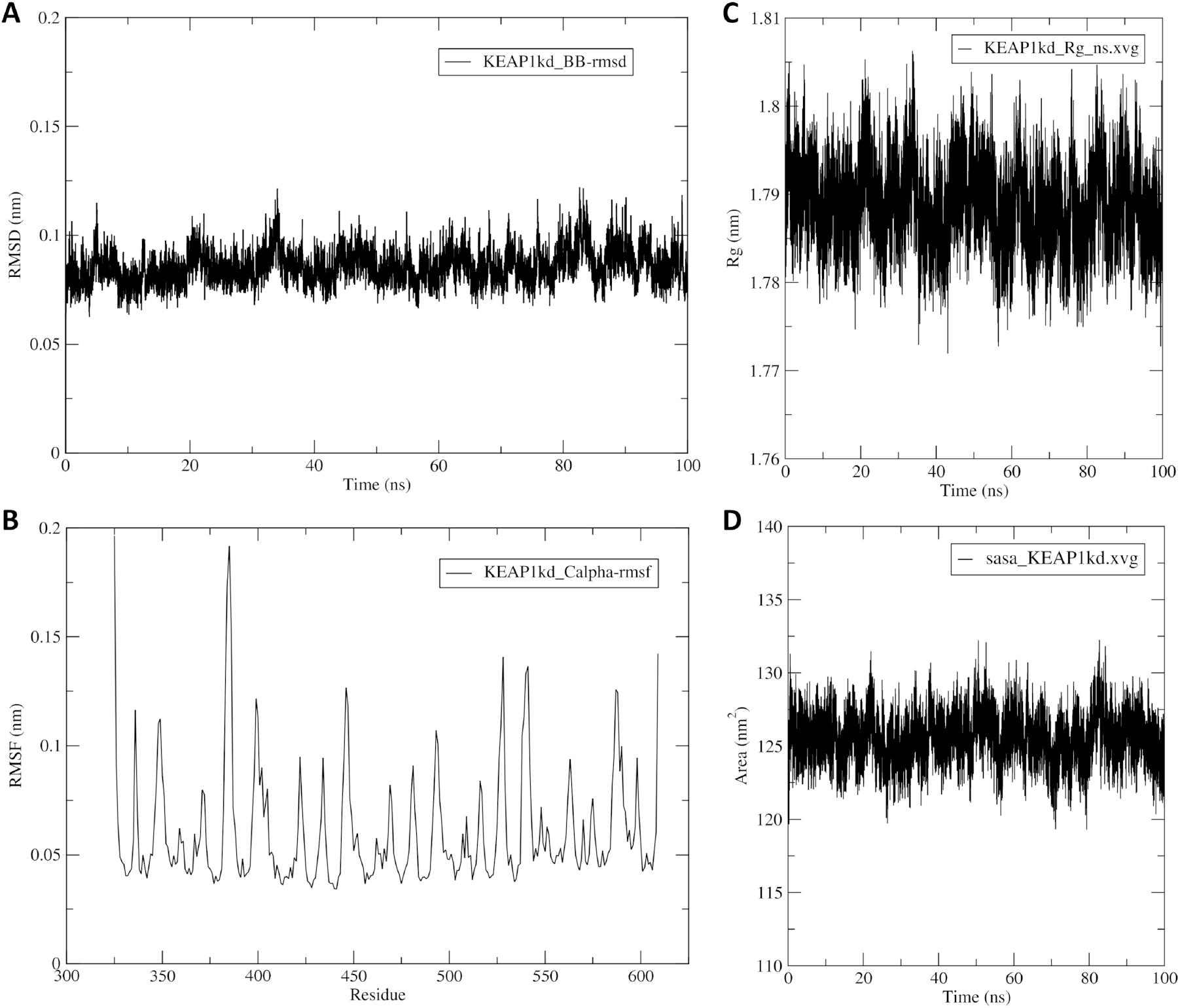
Some trajectory analysis plots generated from the MD simulation of the Kelch domain of the KEAP1 protein (PDB ID 4IQK) as an example. Analyses include the (**A**) Root mean square deviation (RMSD), (**B**) Root mean square fluctuation (RMSF), (**C**) Radius of gyration, and (**D**) Solvent accessible surface area (SASA).

#### 3.2.2 Hydrogen bonding analysis

Hydrogen bond (Hbond) analysis is often used in MD simulations of biomolecules for the investigation and understanding of protein structure, folding, function, and ligand binding as well as other biomolecular interactions [34]. Hbond calculation in GROMACS is carried out using the *gmx hbond* module. Depending on the type of system, the numbers of intra- and inter-molecular Hbonds are calculated and plotted as a function of simulation time. Several other output files, such as the Hbond matrix and index files, are also generated and processed by CHAPERON*g* to parse them as input to other analyses, like the MD Davis-based interactive Hbond matrix calculations.

#### 3.2.3 Principal component analysis

Principal component analysis (PCA) is a statistical technique used to reduce the high-dimensional simulation data–i.e., the coordinates of atoms over time–into a smaller set of orthogonal (principal) components. It helps to visualize the essential dynamics and conformational changes in the trajectory by identifying the most important collective motions in the system [42]. PCA in GROMACS is carried out using the *gmx covar* and *gmx anaeig* modules. The principal components are also processed by CHAPERON*g* and parsed as input for further conformational analyses–e.g., as order parameters for constructing free energy landscapes.

#### 3.2.4 Clustering analysis

Clustering in MD simulation is another common technique that is also used to reduce the complexity of trajectory data. It involves grouping similar conformations based on defined structural similarity, enabling the identification of dominant conformational states, dynamics, and transitions [43]. The *gmx cluster* module in GROMACS carries out the analysis, and the automation by CHAPERON*g* maintains the flexibility and array of options it offers. Figure 2A shows examples of two of the output data plots generated by the analysis.

**Figure 2:**
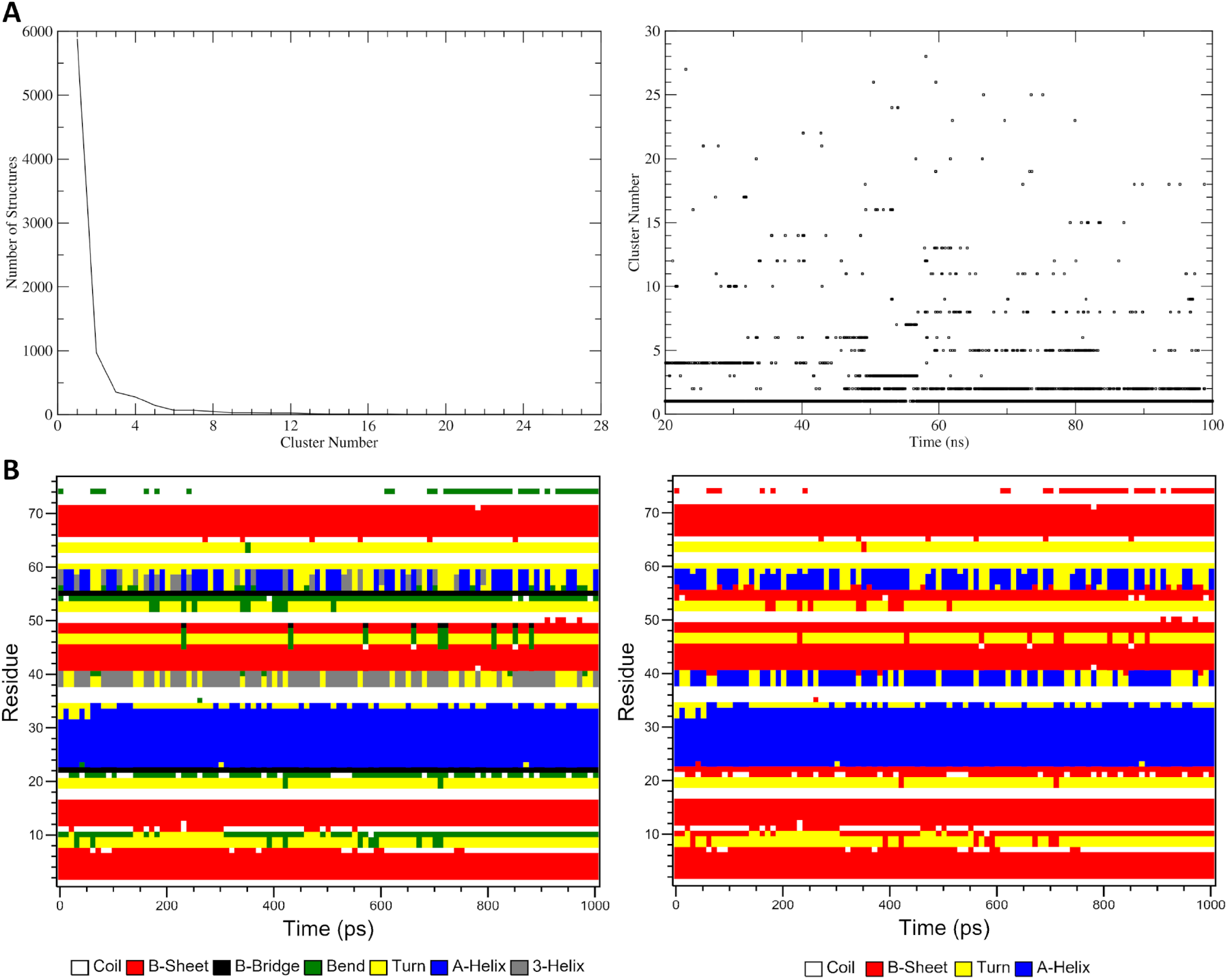
Example output plots of clustering and secondary structure (SS) analyses of MD simulation trajectory. (**A**) Sizes of clusters (*left*) and time-dependent distribution of cluster members (*right*) for the clustering analysis of the ligand-bound Kelch domain of the KEAP1 protein (PDB ID 4IQK) MD simulation trajectory. (**B**) Secondary structure analysis of the human erythrocytic ubiquitin (PDB ID 4GD6) simulation trajectory. Two plots with the seven-SS-type (*left*) and the four-SS-type (*right*) representations are produced.

#### 3.2.5 Secondary structure analysis

Secondary structure (SS) analysis of MD simulation trajectory involves identifying and quantifying the protein secondary structure elements throughout the simulation. The SS analysis in GROMACS is carried out by the *gmx do_dssp* module which relies on the DSSP program [23] for the assignment of SS elements. In addition to the SS analysis plot featuring the default seven SS types assigned by DSSP (see Figure 2B, *left*), CHAPERON*g* reprocesses the SS elements matrix data to generate a second copy of the plot containing only the four basic SS elements—helices, beta-sheets, turns and coils–as shown in Figure 2B (*right*). This simplifies the appearance of the plot to aid its visualization and analysis.

#### 3.2.6 Simulation movie

An MD simulation can be summarized into a movie, which is a collection of frames extracted at a specified interval from the trajectory. Simulation movies facilitate the analysis, interpretation, and communication of the simulation results [44]. They provide an animated overview and visual representation of simulations, and can help to easily visualize the motions of regions of interest, such as active sites and pockets, or to observe conformational movements, interactions or displacement of ligands. The minimum requirement for CHAPERON*g* to create a simulation movie is PyMOL, a widely used molecular visualizer. CHAPERON*g* also utilizes either of the ImageMagick *convert* tool or ffmpeg (when either of them is detected on the user’s machine) for improved movie quality. Supplementary File S1 and Supplementary File S2 show two example movies generated by CHAPERON*g* for the example simulations of ubiquitin and ligand-bound KEAP1 Kelch domain, respectively.

#### 3.2.7 Free energy landscapes

Free energy landscapes (FELs) provide insights into the energetics and stability of different conformational states in MD simulation trajectories. CHAPERON*g* offers three alternative automated ways for the construction of two-dimensional representations of the FEL (Figure 3). These are enabled by the GROMACS *gmx sham* module for 2D visualizations (Figure 3A), the CHAPERON*g* energetic landscape module for enhanced 2D visualizations (Figure 3B), and the MD DaVis tool for interactive 3D visualizations (Figure 3C). Each of these alternatives requires the user to specify two order parameters for the FEL calculations. Global parameters that describe the state of the system can be used as input, including RMSD, Rg, principal components, fraction of native contacts or number of Hbonds, backbone dihedral angles and configurational distance, etc. [45]. Two preset pairs of order parameters–principal components from a PCA run and the RMSD-Rg pair–are available in CHAPERON*g*. A third option that allows the user to provide other quantities of interest as input is also available.

**Figure 3:**
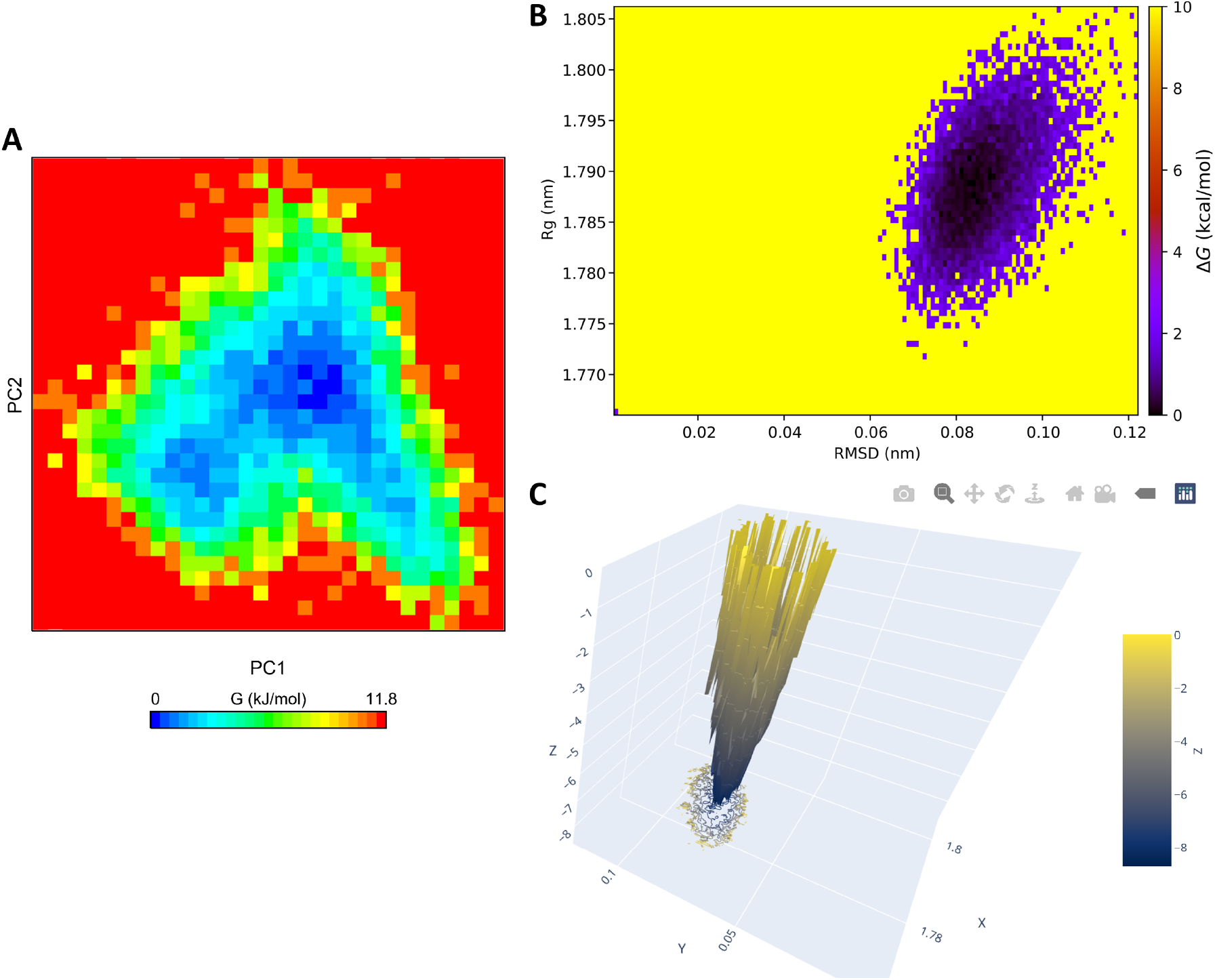
Examples plots of the free energy landscapes (FELs) generated using the free Kelch domain of the KEAP1 protein (PDB ID 4IQK) MD simulation trajectory as an example. (*A*) A 2D plot of the FEL based on principal components generated with gmx sham. (*B*) A CHAPERON*g* -based enhanced 2D plot of the FEL using Rg and RMSD as order parameters. (*C*) An interactive 3D visualization of the Rg-RMSD FEL generated with MD DaVis.

The FEL calculation by CHAPERON*g* employs a modified version of a previously described method [45, 46]. The relative free energies of states are estimated using Boltzmann inversion as shown in Equation 1. The relative free energy of the most probable state is set to zero while other states are computed to have more positive relative free energies. For all the three approaches, CHAPERON*g* also automates the extraction of the lowest energy structures from the FELs, as well as other FEL-guided structures or frames specified by the user.

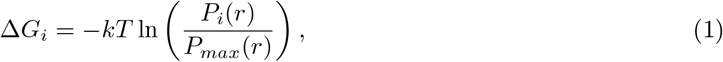

where *k* is the Boltzmann constant, *T* is the simulation temperature, *P*_*i*_(*r*) is the probability of the system being in a particular state *i* characterized by some reaction coordinate *r* (quantities of interest) and is obtained from a histogram of the MD data, *P*_*max*_(*r*) is the probability of the most populated bin (i.e., most probable state), and Δ*G*_*i*_ is the free energy change of the state *i*.

#### 3.2.8 Kernel density estimation

Kernel density estimation (KDE) is a non-parametric technique used to estimate the probability density function (PDF) of a given dataset. This technique utilizes a smooth function, the kernel, centered at sampled datapoints or bins. The Gaussian kernel is one of the commonly used kernels. Given a sample *x* = *x*_1_, *x*_2_, …, *x*_*n*_ with an unknown density f at any given point x. The kernel density estimator of the shape of the function f is defined as shown in Equation 2.

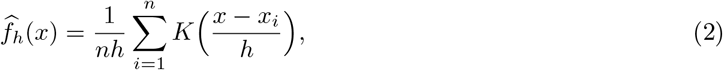

where *K* is the kernel (a simple non-negative function such as the Gaussian distribution), and *h*(*>* 0) is the smoothing bandwidth.

CHAPERON*g* automates the kernel density estimation of the PDF for four common MD trajectory data types, including RMSD, Rg, Hbond, and SASA. This estimation can be carried out for single dataset KDE plots (Figure 4A) as well as for comparative multiple datasets plots (Figure 4B). The plots are generated as .*xvg* and high-quality .*png* files. Depending on the user’s choice, CHAPERON*g* offers automatic and custom selection of the type of bin size estimator, optimal number of histogram bins, and the smoothing bandwidth. The KDE analysis presents a means to gaining further insights into MD simulation trajectories. For instance, the SASA KDE plots shown in Figure 4 provide a different perspective towards the understanding of the SASA data other than the time-dependent information provided in Figure 1D.

**Figure 4:**
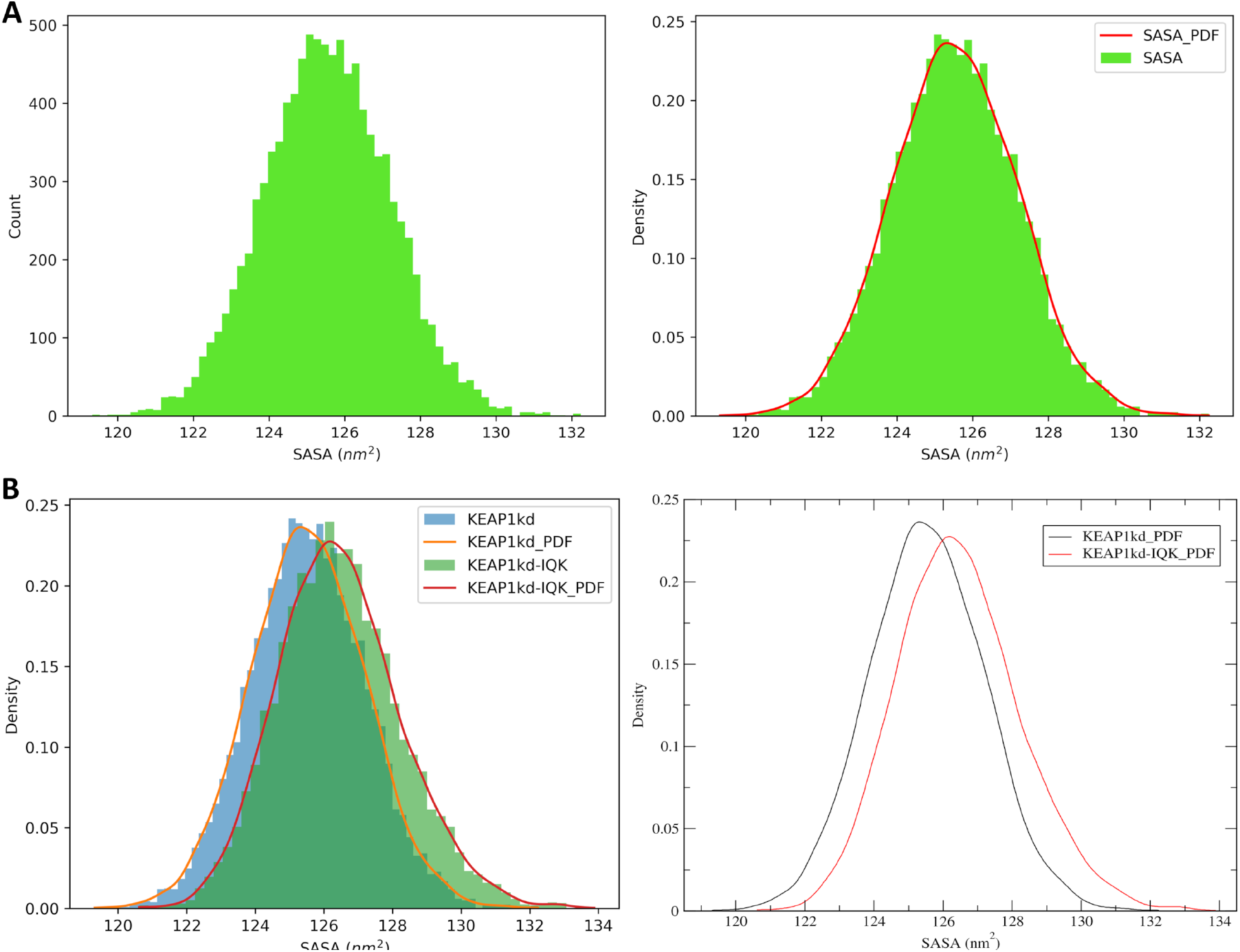
Example CHAPERON*g* kernel density estimation (KDE) plots. (*A*) Histogram (*left*) and KDE (*right*) plots of the SASA data from the example MD simulation trajectory of the KEAP1 Kelch domain. (*B*) Comparative KDE plots of the free KEAP1 Kelch domain protein and the ligand-bound form. Plots are generated as .*png* (*left*) and .*xvg* (*right*) files.

#### 3.2.9 Interactive hydrogen bond matrix

CHAPERON*g* automates the integration of the MD DaVis tool with GROMACS for the construction of a Hbond matrix. To achieve this, CHAPERON*g* prepares a reference structure (from the trajectory) and the Hbond list (from the Hbond index file produced by *gmx hbond*). These files are then parsed as input to MD DaVis to produce an interactive .*html* plot (Figure 5) that gives detailed information about the Hbond contacts recorded in the trajectory [11].

**Figure 5:**
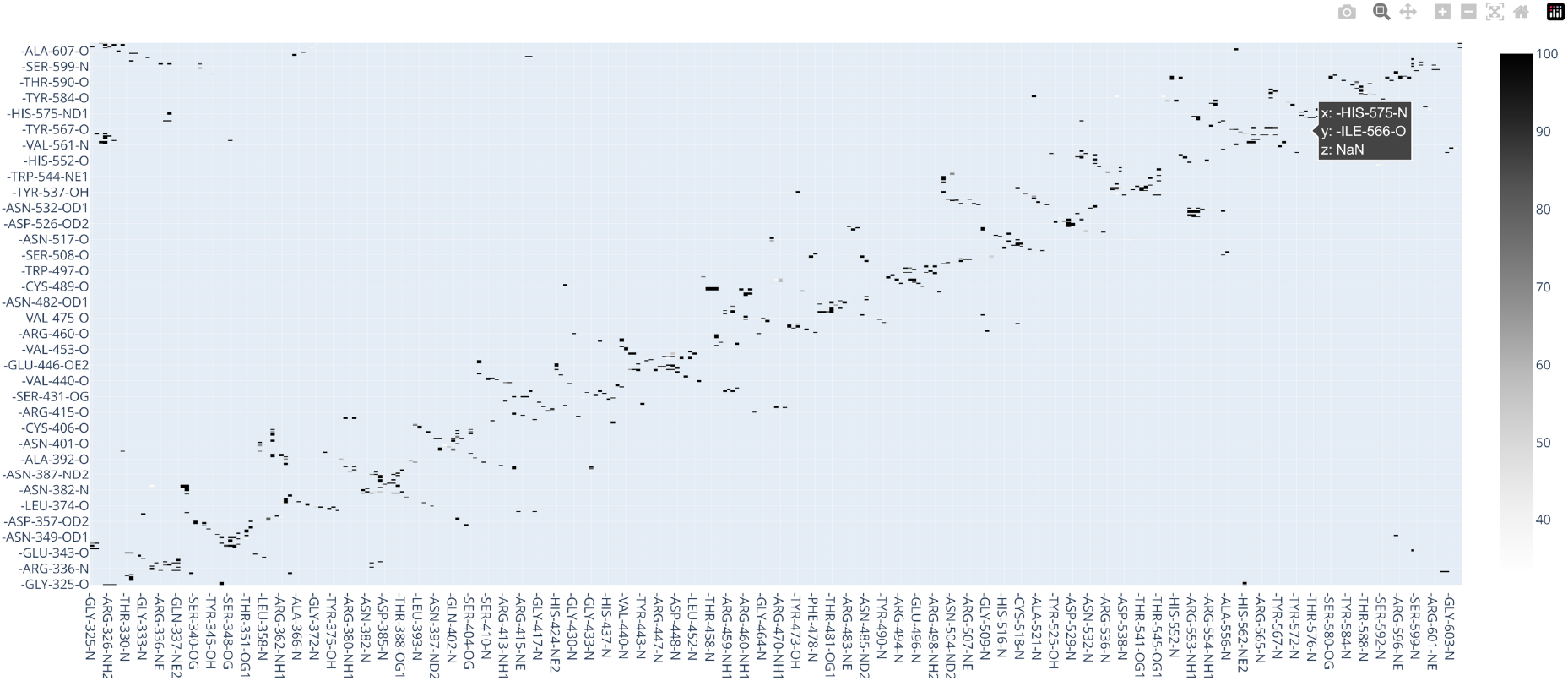
An interactive hydrogen bond matrix generated with MD DaVis using the KEAP1 Kelch domain MD simulation trajectory as an example.

#### 3.2.10 Binding free energy calculations using g_mmpbsa

The g_mmpbsa [47] is a widely used tool that integrates MD simulation with MM-PBSA binding free energy calculation for protein complexes. It also carries out the decomposition of the calculated free energies into contributions per residue [47]. CHAPERON*g* automates and streamlines the workflow for these calculations. Since the original g_mmpbsa is only compatible with GROMACS versions 5.x (or lower) and does not support the more recent and upgraded versions, it’s become a common practice for users to install the older GROMACS version as a second copy for use by g_mmpbsa. Thus, the user would need to provide CHAPERON*g* with the path to the appropriate *gmx* executable. However, the g_mmpbsa code has recently been updated by other people [48] and is supposed to support newer GROMACS version. In this case, there would be no need to specify any path and CHAPERON*g* would automatically call the active GROMACS.

#### 3.2.11 Averaged plot of replica analysis plots

It is a common practice to conduct replica MD simulations of a system, yielding multiple independent trajectories with a higher probability of a wider sampling of the conformational space. Typically, the analysis of the simulations is carried out as means of the replica runs to obtain statistically reliable data, ensure reproducibility, and provide error estimates [49]. CHAPERONg offers a way to automatically generate averaged plots of multiple replica analysis plots, such as the replica plots of the RMSD, Rg, RMSF, SASA, number of hydrogen bonds, or some other user-provided replica plots.

### 3.3 Steered MD and umbrella sampling simulations

The steered MD-umbrella sampling simulation workflow is a powerful technique for estimating the free energy of binding for protein complexes [50, 26, 51], and for studying ligand unbinding pathways [27, 52]. This involves a pulling simulation driven by a biasing potential along a given reaction coordinate. Umbrella sampling simulations are then carried out on a series of configurations in different sampling windows. A technique such as the WHAM is finally used to de-bias the system, calculate the potential of mean force (PMF), and consequently, estimate the free energy of binding. This entire workflow is streamlined and automated by CHAPERON*g* as briefly described below.

#### 3.3.1 System preparation

Depending on the type of system being simulated, the protein and ligand topologies are generated. Then, a placeholder cubic unit cell is generated and the user is interactively guided to adjust the box and center-of-mass dimensions in an iterative visualize-and-adjust manner. This is followed by the solvation, ion adding, energy minimization, and equilibration steps. Several system setup quality assurance analyses are then carried out.

#### 3.3.2 Steered MD simulation and movie

Steered MD simulation involves the pulling apart of the defined pulled and reference groups (illustrated in Figure 6A). Examples of some of the output files are shown in Figure 6, including plots of each of the displacement of the pulled group and the pull force against time (Figures 6B and 6C) and a plot of the pull force against the displacement (Figure 6D). In addition, a movie of the pulling simulation is also generated. Supplementary File S3 and Supplementary File S4 show example movies for protein-protein and protein-ligand steered MD simulations, respectively. Using the PyMOL interface, the user can customize the renderings in the movie, and then re-run CHAPERON*g* to effect the modifications.

**Figure 6:**
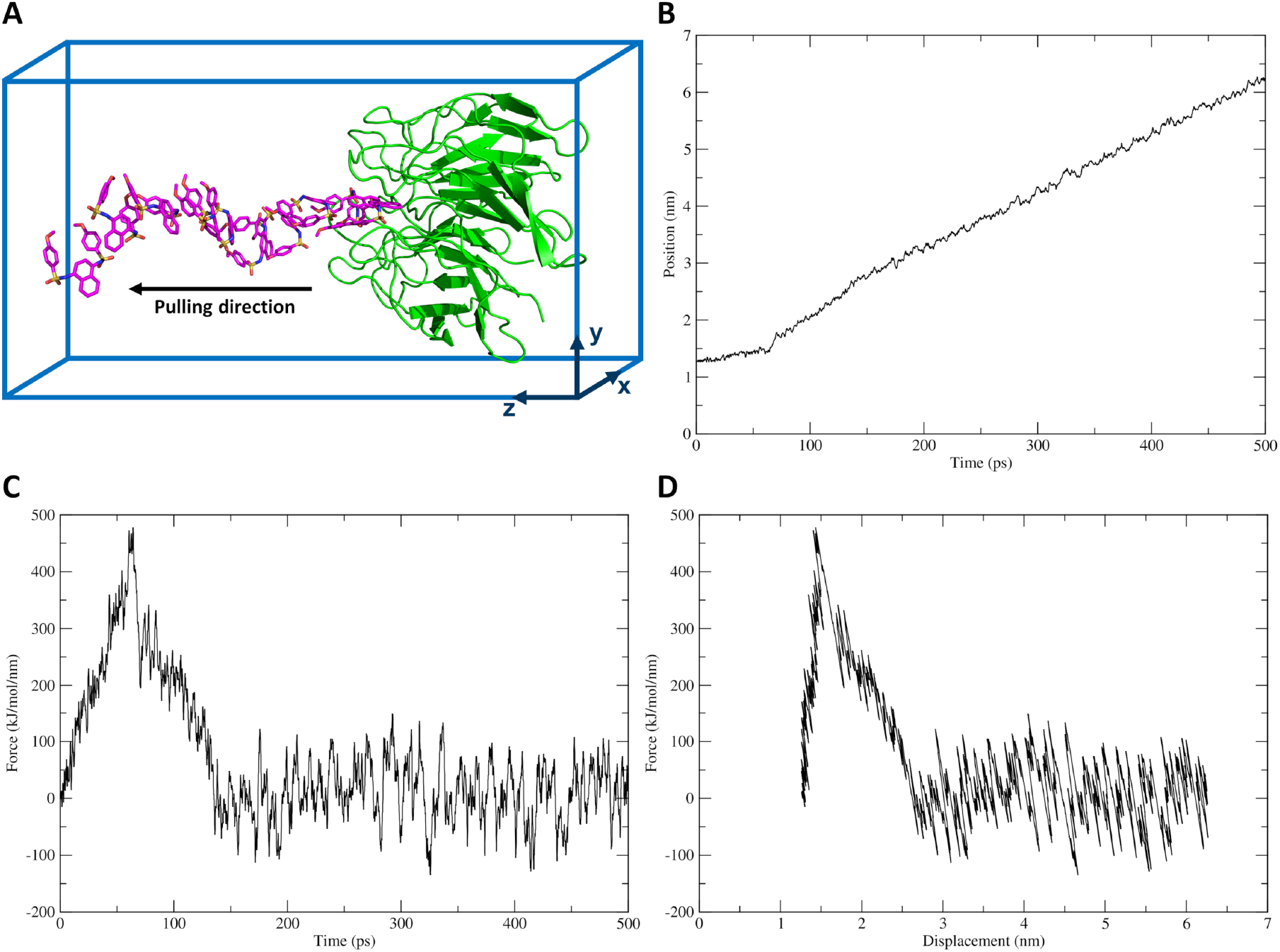
Example steered MD simulation pulling a ligand way from the KEAP1 Kelch domain. (*A*) Illustration of the pulling simulation. The pulled group (ligand) is shown in magenta sticks, and the restrained reference group (receptor) is shown in green cartoon. (*B-D*) Plots of (*B*) pull force against simulation time, (*C*) displacement of the pulled group (the ligand) with time, and (*D*) pull force against the displacement of the pulled group.

#### 3.3.3 Umbrella sampling

Coordinates are extracted from the steered MD trajectory and the COM distance for each frame is calculated using the *gmx distance* module. Based on user-specified spacing, CHAPERON*g* further uses the distances to identify the starting configurations for the umbrella sampling simulations. Umbrella sampling is then iteratively run for each sampling window.

#### 3.3.4 Potential of mean force and binding energy calculation

Using the WHAM calculations via the *gmx wham* module, the output files from the umbrella sampling simulations are used to compute the PMF and, consequently, the free energy of binding. The plots of the umbrella sampling histograms (Figure 7A) and the PMF curve (Figure 7B) are generated as .*png* and .*xvg* files. Also, the binding free energy is calculated and written to a summary file. In a situation where there are windows with insufficient sampling, CHAPERON*g* also offers the possibility to run umbrella sampling for additional user-defined windows.

**Figure 7:**
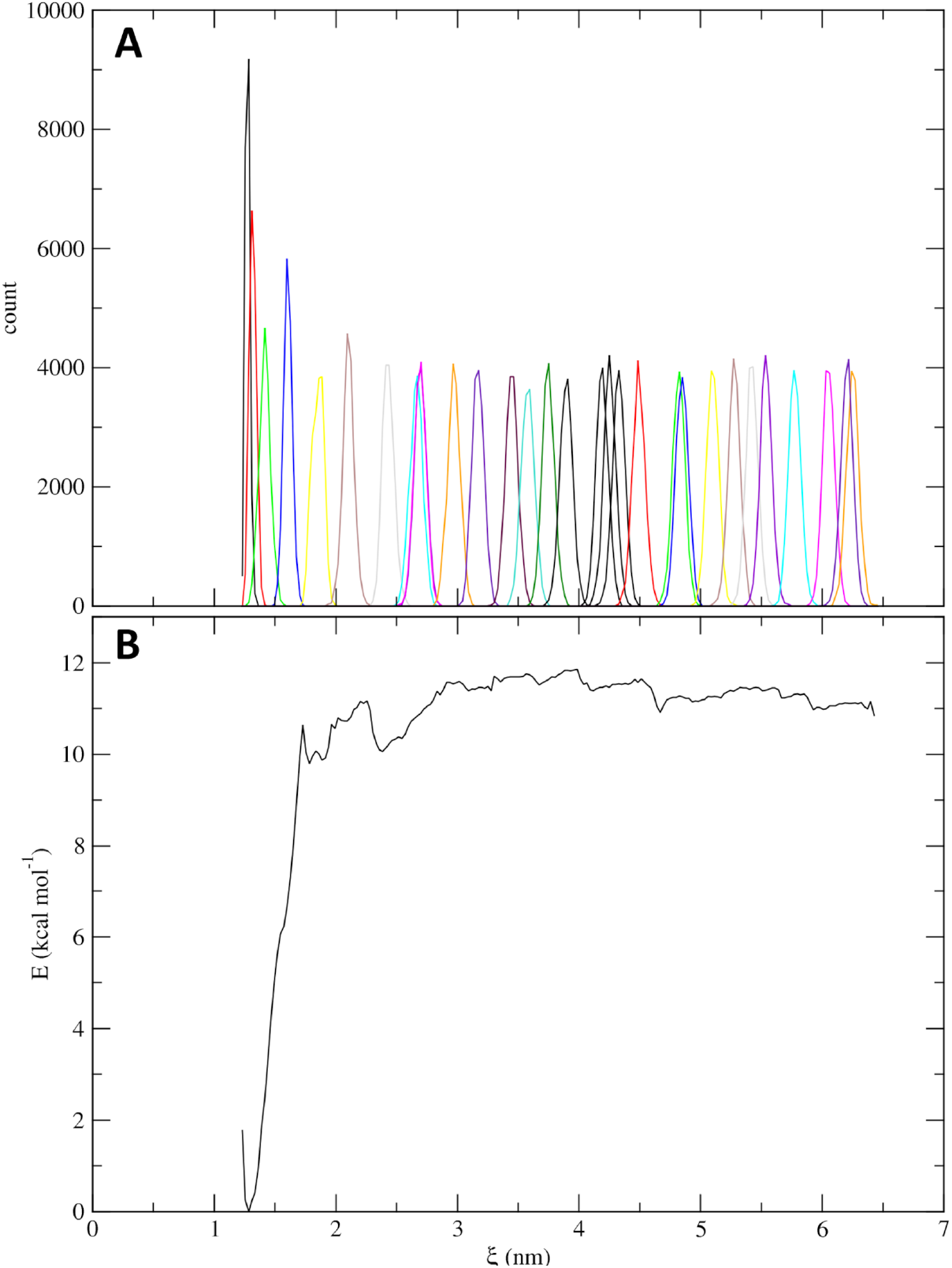
Analysis of an example steered MD and umbrella sampling simulations of the ligand-bound KEAP1 Kelch domain. (**A**) Histograms of the umbrella sampling simulations. (**B**) Potential of mean force (PMF) curve of the ligand unbinding obtained via WHAM calculations.

## 4 Conclusions

In this work, we have developed CHAPERON*g*, an easy-to-use open-source software tool that automates the GROMACS MD simulation pipelines for conventional unbiased MD, steered MD, and enhanced umbrella sampling simulations for diverse biomolecular systems. The tool also offers automated extensive system setup, post-simulation quality assurance analyses, and comprehensive trajectory analyses. Thus, CHAPERON*g* makes MD simulation more accessible to users who have limited experience working with the command line or lack programming skills. It also enables users to gain more insights into MD simulation data by providing an interface to overcome the technical barriers to processing and analyzing trajectory data. We aim to continuously enhance the usability of CHAPERON*g* based on users’ feedback. Future updates would also include additional functionalities to expand the capabilities of the software.

## Notes

### Competing Interest Statement

The authors have declared no competing interest.

## References

[1] Martin Karplus and J Andrew McCammon. Molecular dynamics simulations of biomolecules. Nature Structural Biology, 9(9):646–652, 2002.

[2] Scott A Hollingsworth and Ron O Dror. Molecular dynamics simulation for all. Neuron, 99(6):1129–1143, 2018.

[3] Adam Hospital, Josep Ramon Goñi, Modesto Orozco, and Josep L Gelpí. Molecular dynamics simulations: advances and applications. Advances and Applications in Bioinformatics and Chemistry, pages 37–47, 2015.

[4] David Van Der Spoel, Erik Lindahl, Berk Hess, Gerrit Groenhof, Alan E Mark, and Herman JC Berendsen. GROMACS: fast, flexible, and free. Journal of Computational Chemistry, 26(16):1701–1718, 2005.

[5] Tomasz Makarewicz and Rajmund Kaźmierkiewicz. Molecular dynamics simulation by GROMACS using GUI plugin for PyMOL. Journal of Chemical Information and Modeling, 53(5):1229, 2013.

[6] Arkadeep Sarkar, Jacopo Santoro, Luigi Di Biasi, Francesco Marrafino, and Stefano Piotto. YAMACS: a graphical interface for GROMACS. Bioinformatics, 38(19):4645–4646, 2022.

[7] Hui Liu, Ye Jin, and Hanjing Ding. MDBuilder: a PyMOL plugin for the preparation of molecular dynamics simulations. Briefings in Bioinformatics, 24(2):bbad057, 2023.

[8] Ivo Henrique Provensi Vieira, Eduardo Buganemi Botelho, Thales Junior de Souza Gomes, Roger Kist, Rafael Andrade Caceres, and Fernando Berton Zanchi. Visual dynamics: a web application for molecular dynamics simulation using GROMACS. BMC Bioinformatics, 24(1):1–8, 2023.

[9] Genís Bayarri, Pau Andrio, Adam Hospital, Modesto Orozco, and Josep Lluís Gelpí. BioExcel Building Blocks Workflows (BioBB-Wfs), an integrated web-based platform for biomolecular simulations. Nucleic Acids Research, 50(W1):W99–W107, 2022.

[10] Luciano Porto Kagami, Gustavo Machado das Neves, Luís Fernando Saraiva Macedo Timmers, Rafael Andrade Caceres, and Vera Lucia Eifler-Lima. Geo-Measures: A PyMOL plugin for protein structure ensembles analysis. Computational Biology and Chemistry, 87:107322, 2020.

[11] Dibyajyoti Maity and Debnath Pal. MD DaVis: interactive data visualization of protein molecular dynamics. Bioinformatics, 38(12):3299–3301, 2022.

[12] Pradeep Kota. GUIMACS-a Java based front end for GROMACS. In silico Biology, 7(1):95, 2007.

[13] Sanjit Roopra, Bernhard Knapp, Ulrich Omasits, and Wolfgang Schreiner. jSimMacs for GROMACS: A Java application for advanced molecular dynamics simulations with remote access capability. Journal of Chemical Information and Modeling, 49(10):2412–2417, 2009.

[14] Diamantis Sellis, Dimitrios Vlachakis, and Metaxia Vlassi. Gromita: A fully integrated graphical user interface to gromacs 4. Bioinformatics and Biology Insights, 3:99–102, 2009.

[15] Tomasz Makarewicz and Rajmund Kaźmierkiewicz. Improvements in GROMACS plugin for PyMOL including implicit solvent simulations and displaying results of pca analysis. Journal of Molecular Modeling, 22(109):1–7, 2016.

[16] Kirill Zinovjev and Marc W Van Der Kamp. Enlighten2: molecular dynamics simulations of proteinligand systems made accessible. Bioinformatics, 36(20):5104–5106, 2020.

[17] Adam Hospital, Pau Andrio, Carles Fenollosa, Damjan Cicin-Sain, Modesto Orozco, and Josep Lluís Gelpí. MDWeb and MDMoby: an integrated web-based platform for molecular dynamics simulations. Bioinformatics, 28(9):1278–1279, 2012.

[18] WebGRO for macromolecular simulations. University of Arkansas for Medical Sciences. Available from https://simlab.uams.edu/. Accessed 5 June 2023.

[19] Charles R Harris, K Jarrod Millman, Stéfan J Van Der Walt, Ralf Gommers, Pauli Virtanen, David Cournapeau, Eric Wieser, Julian Taylor, Sebastian Berg, Nathaniel J Smith, et al. Array programming with NumPy. Nature, 585(7825):357–362, 2020.

[20] Pauli Virtanen, Ralf Gommers, Travis E Oliphant, Matt Haberland, Tyler Reddy, David Cournapeau, Evgeni Burovski, Pearu Peterson, Warren Weckesser, Jonathan Bright, et al. Scipy 1.0: fundamental algorithms for scientific computing in python. Nature methods, 17(3):261–272, 2020.

[21] John D Hunter. Matplotlib: A 2D graphics environment. Computing in Science & Engineering, 9(3):90–95, 2007.

[22] Warren L DeLano. The PyMOL molecular graphics system. http://www.pymol.org/, 2002.

[23] Wolfgang Kabsch and Christian Sander. Dictionary of protein secondary structure: Pattern recognition of hydrogen-bonded and geometrical features. Biopolymers, 22(12):2577–2637, 1983.

[24] Justin A Lemkul. From proteins to perturbed Hamiltonians: a suite of tutorials for the GROMACS-2018 molecular simulation package [article v1. 0]. Living Journal of Computational Molecular Science, 1(1):5068, 2018.

[25] Divine Mensah Sedzro, Mukhtar Oluwaseun Idris, Olanrewaju Ayodeji Durojaye, Abeeb Abiodun Yekeen, Adeola Abraham Fadahunsi, and Suleiman Oluwaseun Alakanse. Identifying potential p53-MDM2 interaction antagonists: An integrated approach of pharmacophore-based virtual screening, interaction fingerprinting, MD simulation and DFT studies. ChemistrySelect, 7(39):e202202380, 2022.

[26] Justin A Lemkul and David R Bevan. Assessing the stability of Alzheimer’s amyloid protofibrils using molecular dynamics. The Journal of Physical Chemistry B, 114(4):1652–1660, 2010.

[27] Huiyong Sun, Sheng Tian, Shunye Zhou, Youyong Li, Dan Li, Lei Xu, Mingyun Shen, Peichen Pan, and Tingjun Hou. Revealing the favorable dissociation pathway of type ii kinase inhibitors via enhanced sampling simulations and two-end-state calculations. Scientific Reports, 5(1):8457, 2015.

[28] Shankar Kumar, John M Rosenberg, Djamal Bouzida, Robert H Swendsen, and Peter A Kollman. The weighted histogram analysis method for free-energy calculations on biomolecules. i. the method. Journal of Computational Chemistry, 13(8):1011–1021, 1992.

[29] Marc Souaille and Benoit Roux. Extension to the weighted histogram analysis method: combining umbrella sampling with free energy calculations. Computer Physics Communications, 135(1):40–57, 2001.

[30] Kenno Vanommeslaeghe, Elizabeth Hatcher, Chayan Acharya, Sibsankar Kundu, Shijun Zhong, Jihyun Shim, Eva Darian, Olgun Guvench, P Lopes, Igor Vorobyov, et al. CHARMM general force field: A force field for drug-like molecules compatible with the CHARMM all-atom additive biological force fields. Journal of Computational Chemistry, 31(4):671–690, 2010.

[31] Alan W Sousa da Silva and Wim F Vranken. ACPYPE–antechamber python parser interface. BMC Research Notes, 5(367), 2012.

[32] Daan MF Van Aalten, R Bywater, John BC Findlay, Manfred Hendlich, Rob WW Hooft, and Gert Vriend. PRODRG, a program for generating molecular topologies and unique molecular descriptors from coordinates of small molecules. Journal of Computer-aided Molecular Design, 10:255–262, 1996.

[33] Leela S Dodda, Israel Cabeza de Vaca, Julian Tirado-Rives, and William L Jorgensen. LigParGen web server: an automatic OPLS-AA parameter generator for organic ligands. Nucleic Acids Research, 45(W1):W331–W336, 2017.

[34] Siti Nor Hasmah Ishak, Sayangku Nor Ariati Mohamad Aris, Khairul Bariyyah Abd Halim, Mohd Shukuri Mohamad Ali, Thean Chor Leow, Nor Hafizah Ahmad Kamarudin, Malihe Masomian, and Raja Noor Zaliha Raja Abd Rahman. Molecular dynamic simulation of space and earth-grown crystal structures of thermostable T1 lipase Geobacillus zalihae revealed a better structure. Molecules, 22(10):1574, 2017.

[35] Leandro Martínez. Automatic identification of mobile and rigid substructures in molecular dynamics simulations and fractional structural fluctuation analysis. PloS one, 10(3):e0119264, 2015.

[36] Karen Sargsyan, Cédric Grauffel, and Carmay Lim. How molecular size impacts RMSD applications in molecular dynamics simulations. Journal of chemical theory and computation, 13(4):1518, 2017.

[37] Mukhtar Oluwaseun Idris, Abeeb Abiodun Yekeen, Oluwaseun Suleiman Alakanse, and Olanrewaju Ayodeji Durojaye. Computer-aided screening for potential TMPRSS2 inhibitors: a combination of pharmacophore modeling, molecular docking and molecular dynamics simulation approaches. Journal of Biomolecular Structure and Dynamics, 39(15):5638–5656, 2021.

[38] M Yu Lobanov, NS Bogatyreva, and OV Galzitskaya. Radius of gyration as an indicator of protein structure compactness. Molecular Biology, 42:623–628, 2008.

[39] Castrense Savojardo, Matteo Manfredi, Pier Luigi Martelli, and Rita Casadio. Solvent accessibility of residues undergoing pathogenic variations in humans: from protein structures to protein sequences. Frontiers in Molecular Biosciences, 7(626363), 2021.

[40] Frank Eisenhaber, Philip Lijnzaad, Patrick Argos, Chris Sander, and Michael Scharf. The double cubic lattice method: Efficient approaches to numerical integration of surface area and volume and to dot surface contouring of molecular assemblies. Journal of Computational Chemistry, 16(3):273–284, 1995.

[41] Andrew Shrake and John A Rupley. Environment and exposure to solvent of protein atoms. lysozyme and insulin. Journal of Molecular biology, 79(2):351–371, 1973.

[42] Charles C. David and Donald J. Jacobs. Principal Component Analysis: A Method for Determining the Essential Dynamics of Proteins, pages 193–226. Humana Press, Totowa, NJ, 2014.

[43] Joshua L Phillips, Michael E Colvin, and Shawn Newsam. Validating clustering of molecular dynamics simulations using polymer models. BMC Bioinformatics, 12(1):1–23, 2011.

[44] Erik Lindahl. Molecular Dynamics Simulations, pages 3–26. Springer New York, New York, NY, 2015.

[45] Ivano Tavernelli, Simona Cotesta, and Ernesto E Di Iorio. Protein dynamics, thermal stability, and free-energy landscapes: a molecular dynamics investigation. Biophysical journal, 85(4):2641–2649, 2003.

[46] Elena Papaleo, Paolo Mereghetti, Piercarlo Fantucci, Rita Grandori, and Luca De Gioia. Freeenergy landscape, principal component analysis, and structural clustering to identify representative conformations from molecular dynamics simulations: The myoglobin case. Journal of Molecular Graphics and Modelling, 27(8):889–899, 2009.

[47] Rashmi Kumari, Rajendra Kumar, Open Source Drug Discovery Consortium, and Andrew Lynn. g_mmpbsa–A GROMACS tool for high-throughput MM-PBSA calculations. Journal of Chemical Information and Modeling, 54(7):1951–1962, 2014.

[48] g_mmpbsa (modified). Available from https://github.com/tildeslu/g_mmpbsa. Accessed 15 June 2023.

[49] Bernhard Knapp, Luis Ospina, and Charlotte M Deane. Avoiding false positive conclusions in molecular simulation: the importance of replicas. Journal of Chemical Theory and Computation, 14(12):6127–6138, 2018.

[50] Benjamin Grupp, Justin A. Lemkul, and Thomas Gronemeyer. An in silico approach to determine inter-subunit affinities in human septin complexes. Cytoskeleton, 2023.

[51] Son Tung Ngo, Khanh B Vu, Le Minh Bui, and Van V Vu. Effective estimation of ligand-binding affinity using biased sampling method. ACS omega, 4(2):3887–3893, 2019.

[52] Nguyen Minh Tam, Trung Hai Nguyen, Vu Thi Ngan, Nguyen Thanh Tung, and Son Tung Ngo. Unbinding ligands from SARS-CoV-2 Mpro via umbrella sampling simulations. Royal Society Open Science, 9(1):211480, 2022.

