## Supplementary figures and images for "CHAPERON*g*: A tool for automated GROMACS-based molecular dynamics simulations and trajectory analyses"

### Supplementary File S1

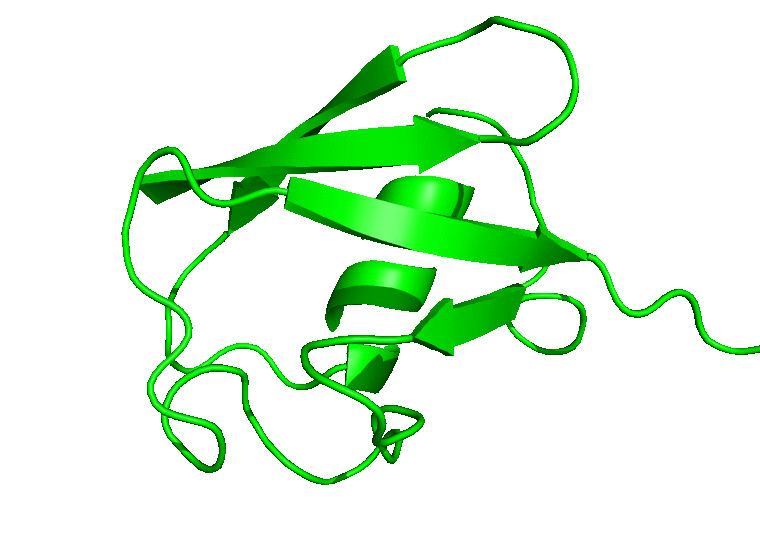

### Supplementary File S2

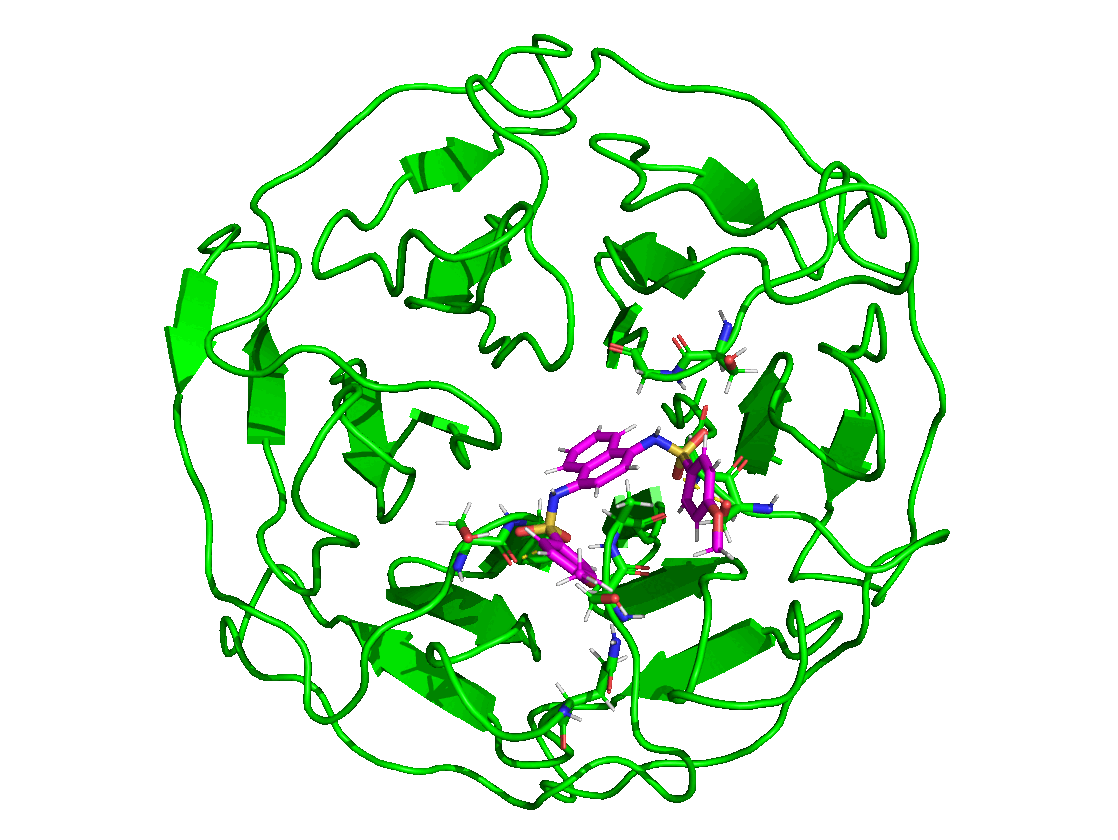

### Supplementary File S3

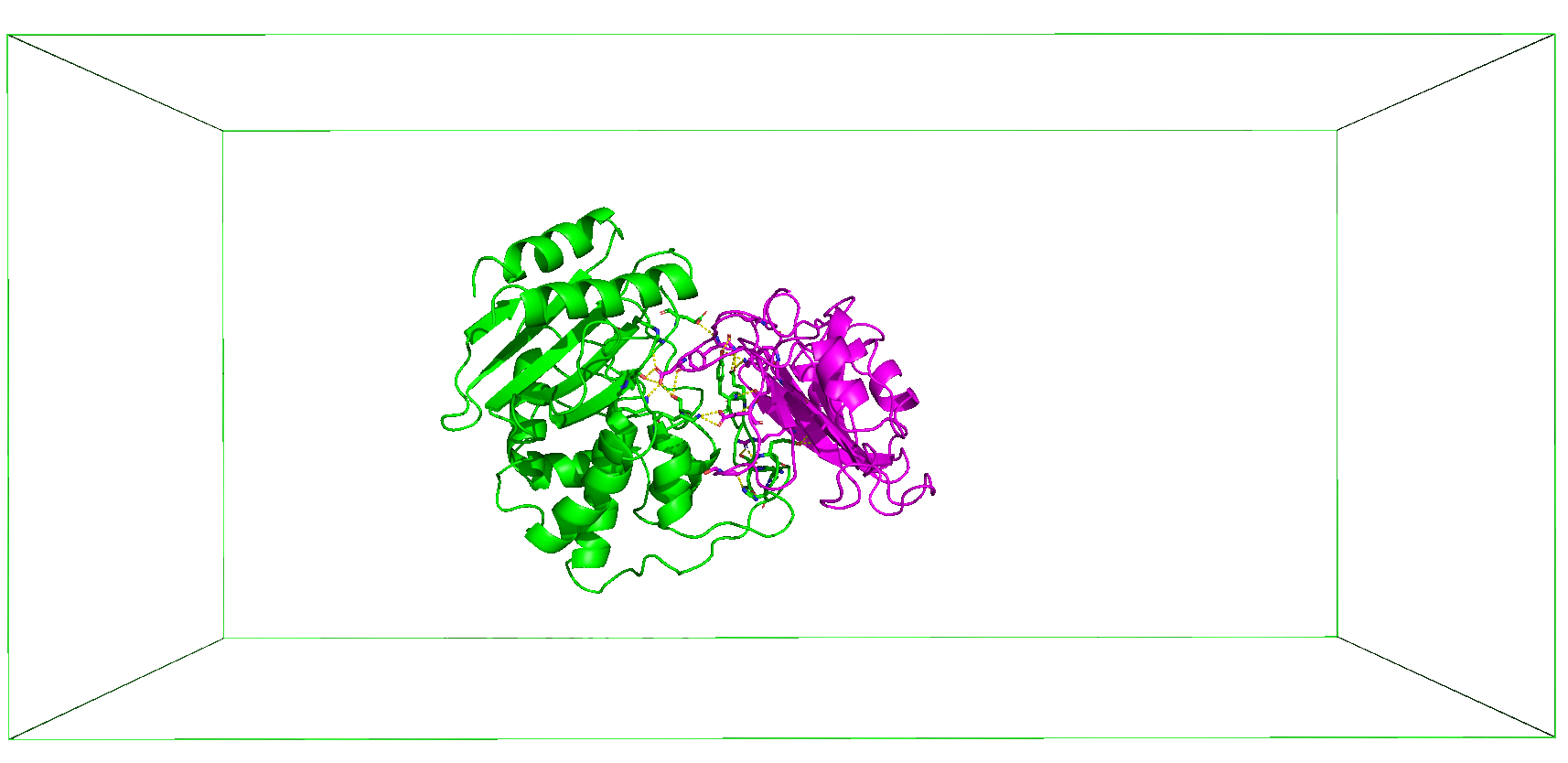

### Supplementary File S4

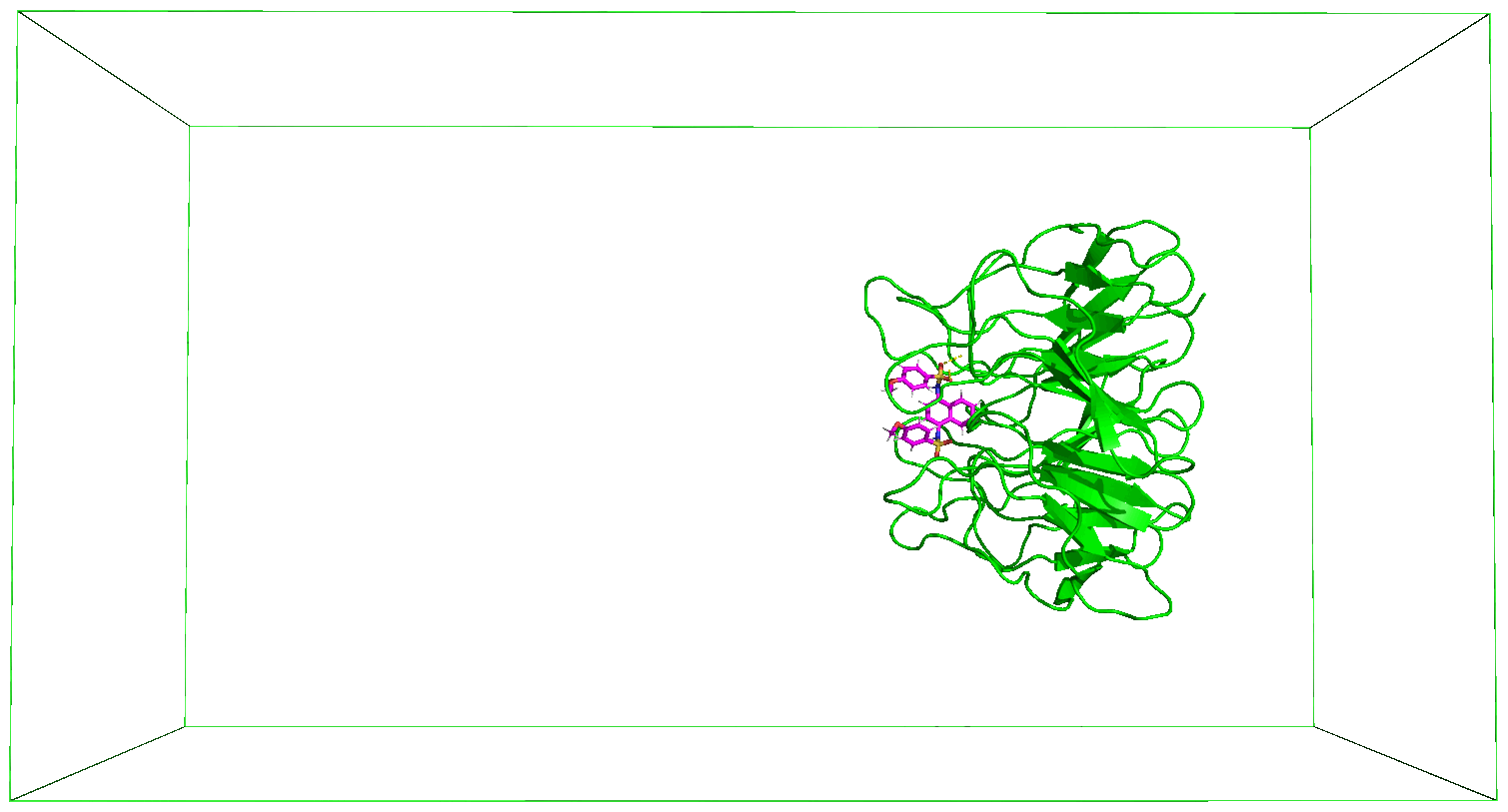
